# NanoDJ: A Dockerized Jupyter Notebook for Interactive Oxford Nanopore MinION Sequence Manipulation and Genome Assembly

**DOI:** 10.1101/586842

**Authors:** Héctor Rodríguez-Pérez, Tamara Hernández-Beeftink, José M. Lorenzo-Salazar, José L. Roda-García, Carlos J. Pérez-González, Marcos Colebrook, Carlos Flores

**Affiliations:** Research Unit, Hospital Universitario Nuestra Señora de Candelaria, Universidad de La Laguna, Santa Cruz de Tenerife, Spain; Genomics Division, Instituto Tecnológico y de Energías Renovables (ITER), Santa Cruz de Tenerife, Spain; Departamento de Ingeniería Informática y de Sistemas, Universidad de La Laguna, Santa Cruz de Tenerife, Spain; Departamento de Matemáticas, Estadística e Investigación Operativa, Universidad de La Laguna, Santa Cruz de Tenerife, Spain; CIBER de Enfermedades Respiratorias, Instituto de Salud Carlos III, Madrid, Spain

**Keywords:** genome analysis, nanopore sequencing, Jupyter, Docker

## Abstract

**Background:** The Oxford Nanopore Technologies (ONT) MinION portable sequencer makes it possible to use cutting-edge genomic technologies in the field and the academic classroom.

**Results:** We present NanoDJ, a Jupyter notebook integration of tools for simplified manipulation and assembly of DNA sequences produced by ONT devices. It integrates basecalling, read trimming and quality control, simulation and plotting routines with a variety of widely used aligners and assemblers, including procedures for hybrid assembly.

**Conclusions:** With the use of Jupyter-facilitated access to self-explanatory contents of applications and the interactive visualization of results, as well as by its distribution into a Docker software container, NanoDJ is aimed to simplify and make more reproducible ONT DNA sequence analysis. The NanoDJ package code, documentation and installation instructions are freely available at https://github.com/genomicsITER/NanoDJ.

## Background

It has never been before so easy and affordable to access and utilize genetic variation of any organism and purpose. This has been motivated by the continuous development of high-throughput DNA sequencing technologies, most commonly known as Next Generation Sequencing (NGS). A key improvement is the possibility of obtaining long single molecule sequences with the fast and cost-efficiency technology released by Oxford Nanopore Technologies (ONT) and the marketing in 2014 of the MinION, a portable, pocket-size, nanopore-based NGS platform [1]. Since then, several algorithms and software tools have flourished specifically for ONT sequence data. Despite its size, it provides multi-kilobase reads with a throughput comparable to other benchtop sequencers in the market (1-10 Gbases by 2017), therefore still necessitating of efficient and integrated bioinformatics tools to facilitate the widespread use of the technology.

While MinION has shown promise in distinct applications [2], because of the low cost, laptop operability, and the USB-powered compact design of MinION, cutting-edge NGS technology is not any more necessarily linked to the established idea of a large machine with high cost that must be located in centralized sequencing centers or in a laboratory bench. As a consequence, the utility of MinION in field experiments to move from sample-to-answers on site have been demonstrated with infectious disease studies [3, 4], off-Earth genome sequencing [5], and species identification in extreme environments [6–8], among others. Leveraging of MinION capabilities in the academic classroom is a natural extension of these field studies to facilitate education of genomics in undergraduate and graduate students [9].

To date, there is no specific software solution aimed to facilitate ONT sequence analyses by integrating capabilities for data manipulation, sequence comparison and assembly in field experiments or for educational purposes to help facilitate learning of genomics [9]. We have developed NanoDJ, an interactive collection of Jupyter notebooks to integrate a variety of software, advanced computer code, and plain contextual explanations. In addition, NanoDJ is distributed as a Docker software container to simplify installation of dependencies and improve the reproducibility of results.

## Features and functionalities

NanoDJ is distributed as a Docker container built underneath Jupyter notebooks, which is increasingly popular in life sciences to significantly facilitate the interactive exploration of data [10], and has been recently integrated in the widely used Galaxy portal [11]. The Docker container allows NanoDJ to run in an isolated, self-contained package, that can be executed seamlessly across a wide range of computing platforms [12], having a negligible impact on the execution performance [13]. NanoDJ integrates diverse applications (Supplementary Table S1) organized into 12 notebooks grouped on three sections (Fig. 1; Table 1). Main results are presented as embedded objects. In addition, one of the notebooks was conceived for educational purposes by setting a particularly simple problem and the inclusion of low-level explanations. To facilitate the use of the educational notebook and bypassing the installation of Docker and NanoDJ, a lightweight version of this notebook and small sets of ONT reads can be utilized from a web-browser using Binder (https://mybinder.org) in the NanoDJ GitHub repository. We illustrate the versatility of NanoDJ in distinct scenarios by providing results from four case studies (Supplementary Text 1).

**Fig. 1.**
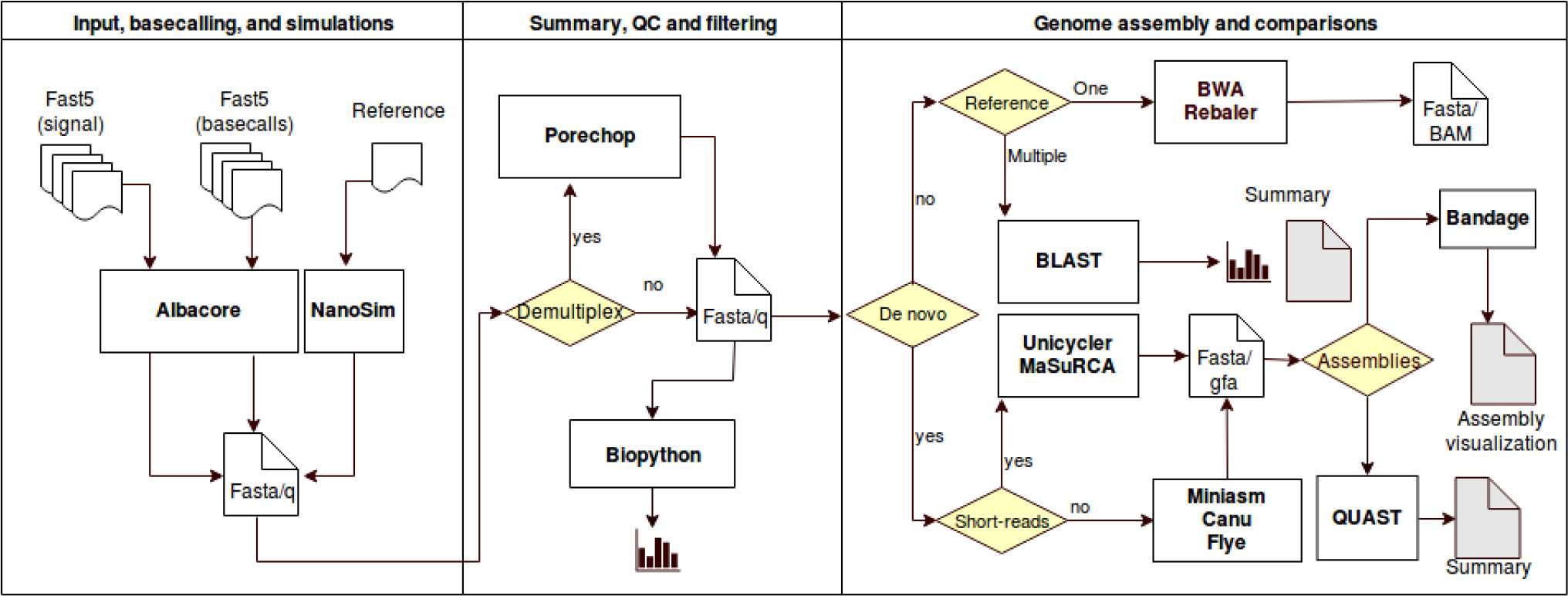
Simplified scheme of all NanoDJ functionalities.

**Table 1.** Summary of NanoDJ notebooks.

| Name | Functionality |
| --- | --- |
| 0.0_QualityControl.ipynb | Evaluate the quality control and sequence handling |
| 1.0_Basecalling.ipynb | Translates the events or the raw electrical signal from an ONT sequencer (FAST5 format) to a DNA sequence to obtain a FASTA or a FASTQ file |
| 1.1_Trim+Demux.ipynb | Perform sequence trimming and demultiplexing |
| 2.0_DeNovo_Canu-Miniasm.ipynb | *De novo* assembly with Canu or Miniasm, and polish with Racon and Pilon |
| 3.0_DeNovo_Canu+polish.ipynb | Nanopolish modules to improve the Canu assembly |
| 4.0_DeNovo_Flye.ipynb | *De novo* assembly with Flye software |
| 5.0_DeNovo_Hybrid.ipynb | Perform *de novo* assembly of Nanopore reads in conjunction with Illumina reads using MaSuRCA and/or Unicycler software |
| 6.0_AssemblyCompare.ipynb | Compare distinct assembly results based on QUAST software |
| 7.0_SimulateReads.ipynb | Obtain simulated reads made with Nanosim software and the Nanosim-h fork with precomputed models |
| 8.0_Alignment.ipynb | Reference-based assembly using either BWA, BLAST or Rebaler software |
| 9.0_AssemblyGraph.ipynb | Assembly graph visualization |
| Educational.ipynb | Performs basecalling (with Albacore), quality control steps, and a BLAST-based classification of the reads (for educational purposes) |

### Input, basecalling, and simulations

Input data can be a list of FAST5 files from previous basecalled runs (e.g. a Metrichor output) or event-level signal data to be basecalled using the latest ONT caller. The user can also simulate reads with NanoSim and pre-computed model parameters. This possibility is important in different scenarios as to help designing an experiment, or to bypass technical difficulties in academic setups [9].

### Summary, quality control and filtering

Either for a simulated or an empirical run, the user will obtain summary data and plots informing of read length distribution, GC content *vs*. length, and read length *vs*. quality score (when available). If barcodes were used in the experiment, Porechop can be used for demultiplexing, barcode trimming and to filter out reads.

### Genome assembly and comparison

Depending on the application, sequence data can be aligned against reference sequences or used for genome assembly using diverse methods. Alignment is performed either against one (BWA and Rebaler) or multiple (BLAST) reference sequences, providing the generation of BAM files for downstream applications (e.g., variant identification) or information of species composition. Alternatively, the user may opt for a *de novo* assembly. NanoDJ allows the use of some of the best-performing algorithms (Canu, Flye, and Miniasm), or to combine ONT reads with others obtained with second-generation NGS platforms for a hybrid assembly (Unicycler and MaSuRCA). The latter provides more effective assemblies and reduced error rate compared to assemblies based only on ONT reads [14]. NanoDJ includes the possibility of contig correction (Racon, Nanopolish, and Pilon). Assemblies can be evaluated with the embedded version of QUAST, and represented with Bandage.

## Limitations and future directions

For non-expert users, it would have been better if NanoDJ was envisaged as an on-line application to facilitate its use. However, our main objective was to integrate major tools for the analysis of ONT sequences in an interactive software environment to facilitate learning the basics behind ONT sequence analysis while providing a useful tool for professionals. Providing it as a Dockerized solution simply bolsters the focus on the use of the tool, reducing the burden of installing all dependencies by the user. At the moment, NanoDJ is set for the analysis of small genomes and targeted NGS studies, although focusing on primary and secondary analysis of DNA sequences. The integration of tools for variant identification and tertiary analysis (annotation of variants or sequence elements, interpretation, etc.) [15–16], as well as for epigenetics [17] and direct RNA sequencing [18] will be the focus of further developments of NanoDJ.

## Conclusions

We present NanoDJ as an integrated Jupyter-based toolbox distributed as a Docker software container to facilitate ONT sequence analysis. NanoDJ is best suited for the analyses of small genomes and targeted NGS studies. We anticipate that the Jupyter notebook-based structure will simplify further developments in other applications.

## Availability and requirements

**Project name**: NanoDJ

**Project home page**: https://github.com/

**Operating system(s):** Windows, Linux, Mac OS

**Programming language**: Bash/Python

**Other requirements**: Docker installation

**License:** GPL

**Any restrictions to use by non-academics**: None

## Aditional files

Table S1: Applications integrated in NanoDJ; **Supplementary Text 1:** Testing on case study datasets (Table S2: Datasets for illustrative uses of NanoDJ; Table S3: Comparison of *de novo* assemblies using different inputs or with an assembly corrector; Table S4: Comparison of three *de novo* assemblers in a high-coverage ONT dataset; Table S5: Comparison of results from two hybrid *de novo* assemblers; Figure S1. Human mitochondrial DNA variant representation against the reference sequence; Table S6: Source of mitochondrial DNA genomes, simulations and classification results.)

## Declarations

### Ethics approval and consent to participate

Not applicable

### Consent for publication

Not applicable

### Availability of data and materials

All data generated or analysed during this study are included in this published article and its supplementary information files.

### Authors’ contributions

HRP scripted and tested the software, and contributed to data analysis; THB was involved in data analysis and interpretation; JLS was involved in data analysis; JRG and CPG revised and tested the software, and revised the manuscript; MC conceived the project, revised and tested the software, and revised the manuscript; CF conceived the project, designed the software, interpreted the data, and critically revised the manuscript.

## Acknowledgements

Not applicable

## Supplementary Material

**Table S1.** Applications integrated in NanoDJ.

| *INPUT, BASECALLING AND SIMULATIONS* |  |  |
| --- | --- | --- |
| *Application* | *Process* | *Result* |
| Albacore<sup>a</sup> v1.0.1 | Basecalling | FASTQ file |
| NanoSim-h<sup>b</sup> v1.0.0.3 | Read simulation | FASTA file with simulated reads |
| *SUMMARY, QUALITY CONTROL AND FILTERING* |  |  |
| *Application* | *Process* | *Result* |
| Biopython<sup>c*</sup> v1.70 | Summarization | Plots and tables |
| Porechop<sup>d</sup> v0.2.2 | Demultiplexing, trimming and filtering | Demultiplexed FASTQ files |
| *GENOME ASSEMBLY AND COMPARISONS* |  |  |
| *Application* | *Process* | *Result* |
| BWA<sup>e</sup> v0.7.17 | Alignment (one reference) | BAM file |
| Rebaler<sup>f</sup> v0.1.0 | Alignment (one reference) | FASTA file |
| BLAST<sup>g</sup> v2.7.1 | Alignment (multiple reference) | Read assignment (summary) |
| Miniasm<sup>h</sup> r104 | De novo assembly | Assembly files (FASTA, PAF) |
| Flye<sup>i</sup> v2.3.1 | De novo assembly | Assembly files (FASTA) |
| Canu<sup>j</sup> v1.6 | De novo assembly | Assembly files (FASTA, .gfa) |
| Racon<sup>k</sup> v0.5.0 | Contig correction | Corrected FASTA |
| Nanopolish<sup>l</sup> v0.8.5 | Contig correction | Corrected FASTA |
| Pilon<sup>m</sup> v1.22 | Contig correction | Corrected FASTA |
| Unicycler<sup>n**</sup> v0.4.1 | Hybrid de novo assembly | Assembly files (FASTA, .gfa) |
| MaSuRCA<sup>o</sup> v3.2.2 | Hybrid de novo assembly | Assembly files (FASTA, .gfa) |
| QUAST<sup>p</sup> v5.0.0 | Assembly comparison | Summary tables and plots |
| Bandage<sup>q</sup> v0.8.1 | Visualization of assembly graphs (.gfa) | Assembly plot |

<sup>a</sup>https://nanoporetech.com/; <sup>b</sup>Yang et al., 2017 (https://pypi.python.org/pypi/NanoSim-H); <sup>c</sup>Cock et al., 2009 (www.biopython.org); <sup>d</sup>https://github.com/rrwick/Porechop; <sup>e</sup>Li & Durbin 2010 (https://sourceforge.net/projects/bio-bwa/files); <sup>f</sup>https://github.com/rrwick/Rebaler; <sup>g</sup>Altschul et al., 1990 (<ftp://ftp.ncbi.nlm.nih.gov/blast/executables/blast+/LATEST>); <sup>h</sup>Lin et al., 2016 (https://github.com/lh3/miniasm); <sup>i</sup>Kolmogorov et al., 2018 (https://github.com/fenderglass/Flye); <sup>j</sup>Koren et al., 2017 (https://github.com/marbl/canu); <sup>k</sup>Vaser et al., 2017 (https://github.com/isovic/racon); <sup>l</sup>https://github.com/jts/nanopolish; <sup>m</sup>Walker et al., 2014 (https://github.com/broadinstitute/pilon); <sup>n</sup>Wick et al., 2017 (https://github.com/rrwick/Unicycler); <sup>o</sup>Zimin et al., 2013 (<ftp://ftp.genome.umd.edu/pub/MaSuRCA>); <sup>p</sup>Gurevich et al., 2013 (https://sourceforge.net/projects/quast); <sup>q</sup>Wick et al., 2015 (https://rrwick.github.io/Bandage/)
\*Dependencies included: Numpy, Matplotlib and Pandas.
\*\*Dependencies included: Spades, Racon, Pilon, bowtie2, samtools, blast+.

### Supplementary Text 1. Testing on case study datasets

We illustrate the versatility of NanoDJ in distinct scenarios with four example datasets starting from diverse file inputs (Supplementary Table S2): (1) the assembly of a bacterial genome testing distinct *de novo* assemblers based on a high-coverage ONT data (different inputs); (2) a hybrid genome assembly of a bacterial genome based on low-pass ONT sequencing and short-read data at high coverage; (3) an emulation of a resequencing experiment to map ONT reads to a reference sequence for the identification of genetic variants (with third-party tools); and (4) an evaluation of species composition based on simulated ONT reads. All analyses were performed in a Ubuntu 16.04 Server with two Intel Xenon E5-2650 12-core 2.2 GHz processors and 256 Gb of RAM.

**Table S2.**
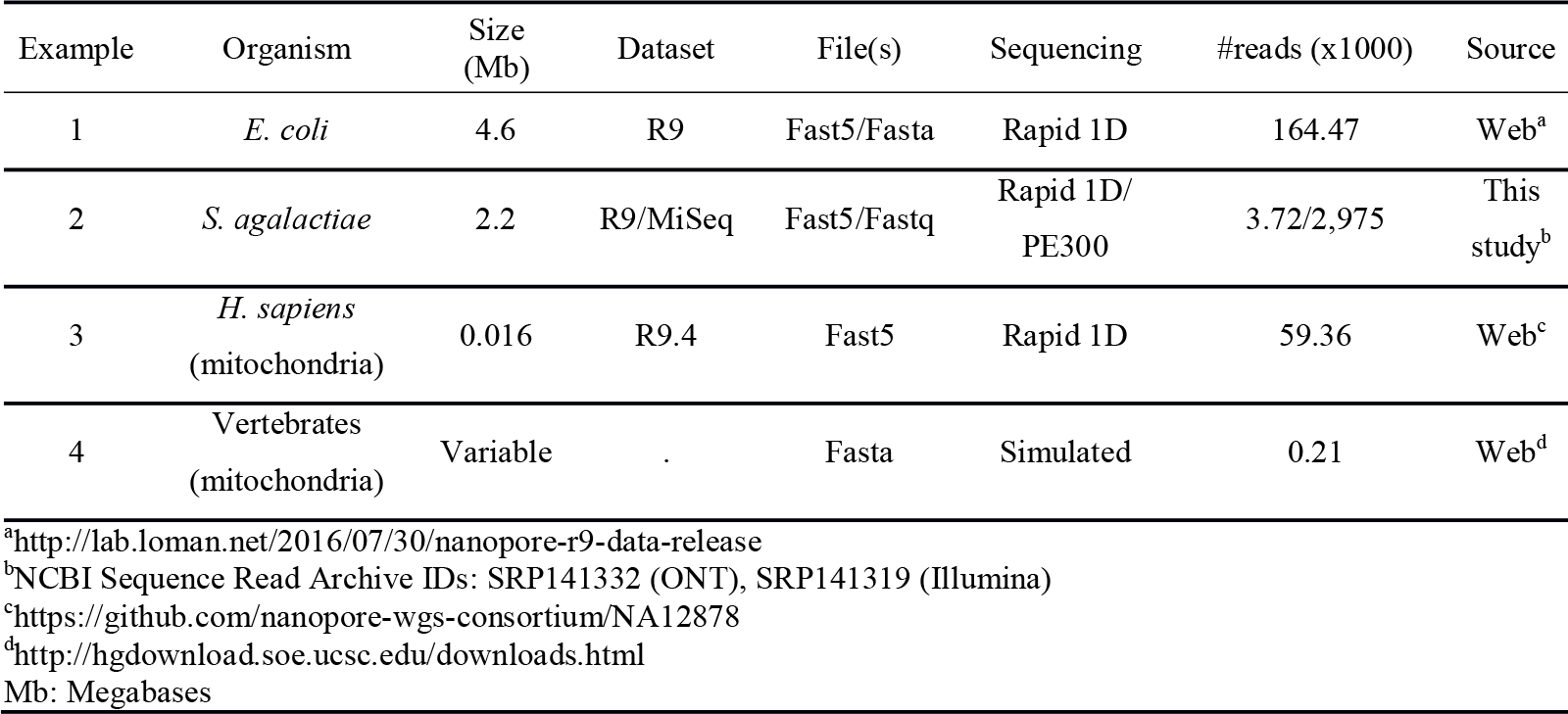
Datasets for illustrative uses of NanoDJ.

#### Example 1.

We used NanoDJ to test different *de novo* assemblers using default parameters in a 4.6 Mb reference *E. coli* genome (K-12-MG1655; NC_000913.3) obtained at a high coverage (>20X) with ONT. For these analyses we have used the following notebooks in the order: 1^st^) 1.0_Basecalling.ipynb, 2^nd^) 2.0_DeNovo_Canu-Miniasm.ipynb, 3^rd^) 4.0_DeNovo_Flye.ipynb, and 4^th^) 6.0_AssemblyCompare.ipynb. A first test with the most accurate assembler (Canu) compared the use of FAST5 files (333 Gb) and of a FASTA (1.5 Gb) file as the input dataset. In both cases, Canu assembled all reads into a single 4.6 Mb genome covering 99.9% of the reference. Using FASTA as the input did not reduced the elapsed time and introduced more mismatches and indels (Supplementary Table S3). The use of assembly polishers (e.g. Racon) did not improve significantly the assembly. Besides, we used the FASTA file to compare three distinct assemblers (Canu, Flye, and Miniasm). Overall, Canu and Flye were by far the most accurate alternatives, but Flye was much less computationally expensive. Miniasm was able to assembly the genome into a single contig, and was the fastest alternative of all three (Supplementary Table S4).

**Table S3.** Comparison of Canu-based *de novo* assemblies of *E. coli* K-12-MG1655^$^ using different inputs or with an assembly corrector.

| Parameters | FASTA | FAST5 | FAST5-Racon |
| --- | --- | --- | --- |
| Total length assembled (bp) | 4,602,643 | 4,662,047 | 4,706,877 |
| Contigs (>=500 bp) | 1 | 1 | 1 |
| Genome fraction (%)<sup>*</sup> | 99.88 | 99.98 | 99.99 |
| Mismatches (#)* | 14,674 | 8,446 | 10,533 |
| Indels (#)* | 55,889 | 24,187 | 21,805 |
| N50 | 4,602,643 | 4,662,047 | 4,706,877 |
| GC (%) | 51.07 | 50.95 | 50.75 |
| Elapsed time (sec.) | 138,221.92 | 15,553.49 | 15,646.13 |

**Table S4.** Comparison of three de novo assemblers in a high-coverage ONT dataset (FASTA input) obtained from E. coli K-12-MG1655^$^.

| Parameters | Canu | Flye | Miniasm |
| --- | --- | --- | --- |
| Total length assembled (bp) | 4,602,643 | 4,678,264 | 4,404,394 |
| Contigs (>=500 bp) | 1 | 1 | 1 |
| Largest alignment (pb)* | 3,336,128 | 2,310,685 | 77 |
| Mismatches (#)* | 14,674 | 12,678 | 0 |
| Indels (#)* | 55,889 | 40,090 | 2 |
| N50 | 4,602,643 | 4,678,264 | 4,404,394 |
| GC (%) | 51.07 | 50.72 | 52.48 |
| Elapsed time (sec.) | 138,221.92 | 1,038 | 104 |

<sup>§</sup>Genome size and GC content are 4,641,652 bp and 50.79%, respectively.
\*Against *E. coli* reference sequence NC\_000913.3.
Indels: insertion/deletion variants; N50: minimum contig length to cover at least 50% of the genome; GC: guanine-cytosine content

#### Example 2.

Here we used available *S. agalactiae* reads from a low-pass (~2.5X) ONT MinION experiment and 2,97 million reads (>300X) from a MiSeq (Illumina, Inc.) run to compare two state-of-the-art hybrid assemblers. In this case, we have used the following notebooks in the order: 1^st^) 1.0_Basecalling.ipynb, 2^nd^) 5.0_DeNovo_Hybrid.ipynb, 3^rd^) 6.0_AssemblyCompare.ipynb, and 4^th^) 9.0_AssemblyGraph.ipynb. NanoDJ integrated basecalling with Albacore, the hybrid assembly with Unicycler or MaSuRCA, and the assembly comparisons. Unicycler was superior to MaSuRCA based on the number of contigs, N50, the size of the largest contig, the number of mismatches and indels, and the proportion of covered genome (Supplementary Table S5). Plotting of Unicycler results also allowed isolating a 2.5 Kb plasmid sequence supported by our previous findings (unpublished), while MaSuRCA split its sequence into two small contigs.

**Table S5.**
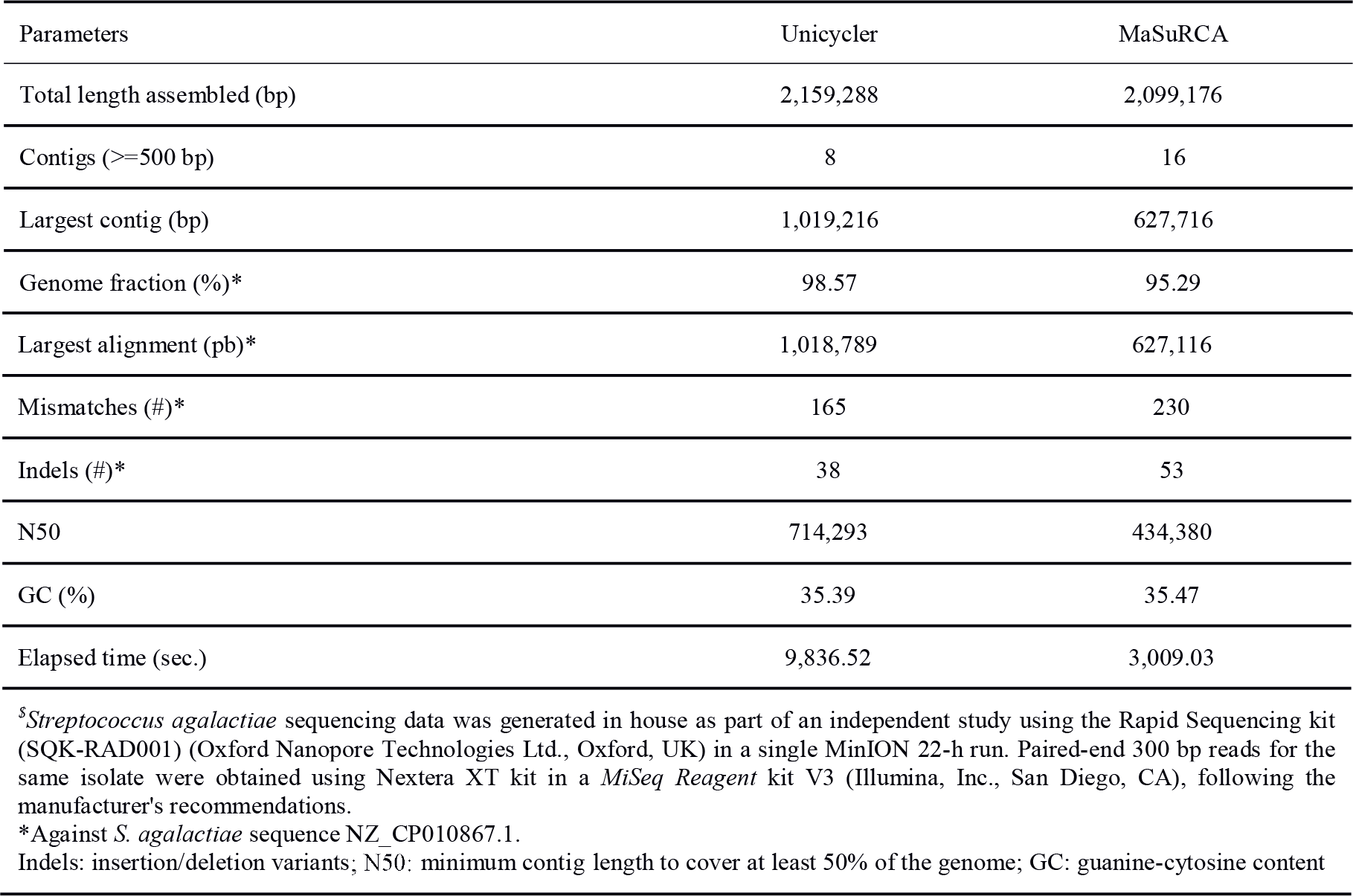
Comparison of results from two hybrid *de novo* assemblers in a *S. agalactiae* dataset^$^.

| Parameters | Unicycler | MaSuRCA |
| --- | --- | --- |
| Total length assembled (bp) | 2,159,288 | 2,099,176 |
| Contigs (>=500 bp) | 8 | 16 |
| Largest contig (bp) | 1,019,216 | 627,716 |
| Genome fraction (%)* | 98.57 | 95.29 |
| Largest alignment (pb)* | 1,018,789 | 627,116 |
| Mismatches (#)* | 165 | 230 |
| Indels (#)* | 38 | 53 |
| N50 | 714,293 | 434,380 |
| GC (%) | 35.39 | 35.47 |
| Elapsed time (sec.) | 9,836.52 | 3,009.03 |

<sup>S</sup>*Streptococcus agalactiae* sequencing data was generated in house as part of an independent study using the Rapid Sequencing kit (SQK-RAD001) (Oxford Nanopore Technologies Ltd., Oxford, UK) in a single MinION 22-h run. Paired-end 300 bp reads for the same isolate were obtained using Nextera XT kit in a *MiSeq Reagent* kit V3 (Illumina, Inc., San Diego, CA), following the manufacturer's recommendations.
\*Against *S. agalactiae* sequence NZ\_CP010867.1.
Indels: insertion/deletion variants; N50: minimum contig length to cover at least 50% of the genome; GC: guanine-cytosine content

#### Example 3.

We used FAST5 files available for reads mapping to the human mitochondrial DNA (33 Gb) from the NA12878 human genome reference standard on the ONT MinION. NanoDJ was used to extract all reads and obtain a FASTQ file, integrate all reads into a single FASTA assembly by mapping against the revised Cambridge Reference Sequence (NC_012920, gi:251831106) with Rebaler, and obtain a BAM file, which was then manually inspected with IGV (Robinson et al., 2011) (Supplementary Fig. S1). In this case study, we have used the following notebooks in the order: 1^st^) 1.0_Basecalling.ipynb, 2^nd^) 0.0_QualityControl.ipynb, and 3^rd^) 8.0_Alignment.ipynb. The assembly was then used in HAPLOFIND, a third-party tool based on PhyloTree build 17 (Vianello et al., 2013), to confirm the classification of the reference data as H13a1a1 mitochondrial haplogroup.

**Figure S1.**
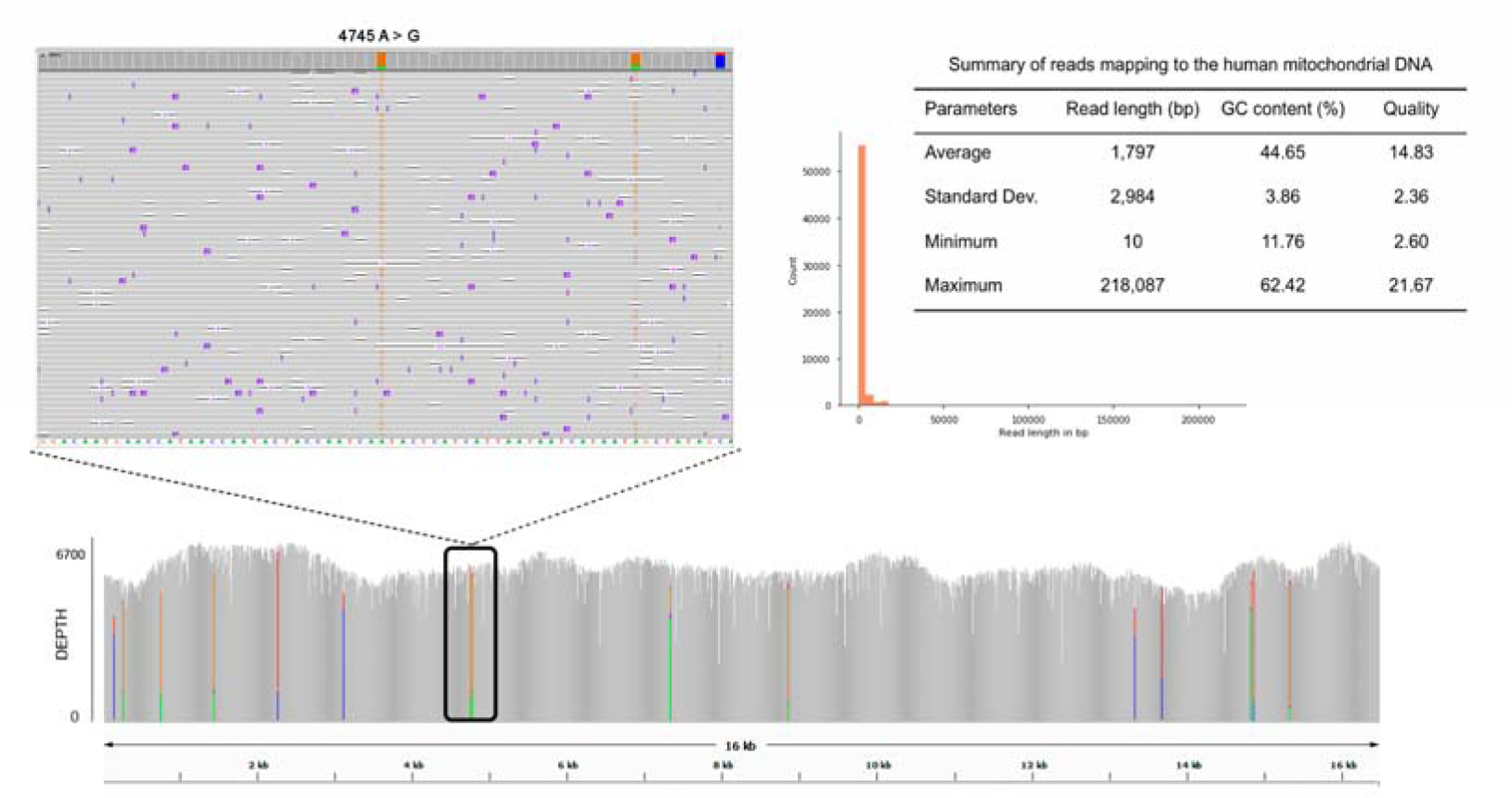
Human mitochondrial DNA variant representation against the reference sequence (left), read distribution (right), and coverage information (bottom). A consensus calling of variant alleles was made if the allele was present in >70% of reads. This identified the correct nucleotide at all reference positions, permitting the identification of the expected haplogroup for the reference DNA. The figure illustrates the case of position 4,745 where a A>G change is supported by the reads, defining H13a1 haplogroup.

#### Example 4.

We used NanoDJ to create a local BLAST database with seven mitochondrial reference genomes from distinct vertebrates. In parallel, NanoDJ was used to simulate 30 ONT reads from seven species with NanoSim-h, which were merged into a single FASTA to emulate a 210-read heterogeneous sample run with balanced proportion of species (~14.3% each). NanoDJ was finally used for a BLAST-based classification of simulated reads to obtain species abundance. In this case study, we have used the following notebooks in the order: 1^st^) 7.0_SimulateReads.ipynb, 2^nd^) 0.0_QualityControl.ipynb, and 3^rd^) 8.0_Alignment.ipynb. As expected, the average abundance was supported by 14.3% of reads (excluding the unassigned), although with minimal fluctuations between 12.9 and 16.1% (Supplementary Table S6).

**Table S6.** Source of mitochondrial DNA genomes, simulations and classification results.

| Species | Reference | Simulated reads | Assigned reads | Proportion* |
| --- | --- | --- | --- | --- |
| *Gallus gallus* | Dec. 2015 (Gallus_gallus-5.0/galGal5) | 30 | 27 | 14.5 |
| *Alligator mississippiensis* | Aug. 2012 (allMis0.2/allMis1) | 30 | 25 | 13.4 |
| *Bos taurus* | Jun. 2014 (UMD_3.1.1/bosTau8) | 30 | 27 | 14.5 |
| *Equus caballus* | Sep. 2007 (Broad/equCab2) | 30 | 29 | 15.6 |
| *Oreochromis niloticus* | Jan. 2011 (Nile tilapia/oreNil2) | 30 | 30 | 16.1 |
| *Rattus norvegicus* | Jul. 2014 (RGSC 6.0/rn6) | 30 | 24 | 12.9 |
| *Ovis aries* | Aug. 2012 (ISGC Oar_v3.1/oviAri3) | 30 | 24 | 12.9 |

\*Excluding 24 reads that were unassigned.

